# Preparation of staged *Caenorhabditis elegans* embryos using size filtration

**DOI:** 10.1101/2025.01.14.632944

**Authors:** Nikita S. Jhaveri, Maya K. Mastronardo, J.B. Collins, Erik C. Andersen

## Abstract

The free-living nematode *Caenorhabditis elegans* has been routinely used to study gene functions, gene interactions, and conserved signaling pathways. Most experiments require that the animals are synchronized to be at the same specific developmental stage. Bleach synchronization is traditionally used to obtain a population of staged embryos, but the method can have harmful effects on the embryos. The physical separation of differently sized animals is preferred but often difficult to perform because some developmental stages are the same sizes as others. Microfluidic device filters have been used as alternatives, but they are expensive and require customization to scale up the preparation of staged animals. Here, we present a protocol for the synchronization of embryos using mesh filters. Using filtration, we obtained a higher yield of embryos per plate than using standard bleach synchronization protocol and at a scale beyond microfluidic devices. Importantly, filtration does not affect downstream larval development assays as much as bleach synchronization does. In conclusion, we have exploited the differences in the sizes of *C. elegans* developmental stages to isolate embryo cultures suitable for use in high-throughput assays.

## Introduction

Nematodes of the genus *Caenorhabditis* are an ideal system to study genes, genetic interactions, and evolutionary conserved signaling pathways. The most studied organism of this group is *Caenorhabditis elegans*, because of its short life cycle, hermaphroditic mating system, cost-effective cultivation, small size, transparency, small genome, and the simplicity of genetic modification and functional testing in a laboratory setting (1). *C. elegans* develop through four morphologically distinct larval stages (L1 - L4), before progressing into adults. Most experiments require the nematodes to be at a specific developmental stage because of the distinct morphologies, behaviors, and gene expression patterns of each stage. High-throughput assays such as RNA sequencing, genome-wide association studies (GWAS), drug screens, proteomics, and behavioral assays require a large number of animals at a particular stage. The standard way of obtaining nematodes at a specific developmental stage is by collecting embryos and allowing them to hatch into L1 larvae that are arrested when starved. The synchronous population of L1 larvae is fed and then incubated at the desired temperature until the animals grow to the target developmental stage. Embryos are obtained from the uterus of gravid adults by dissolving the hypodermis using a hypochlorite (bleach) solution (Figure 1). However, different parameters such as the bleaching time and temperature of the bleach solution affect the survival of the embryos (2). To control for the variability in the numbers of viable, unaffected animals after bleach synchronization, three independent replicates are typically performed for high-throughput assays, making the process time-consuming (3–5).

**Figure 1.**
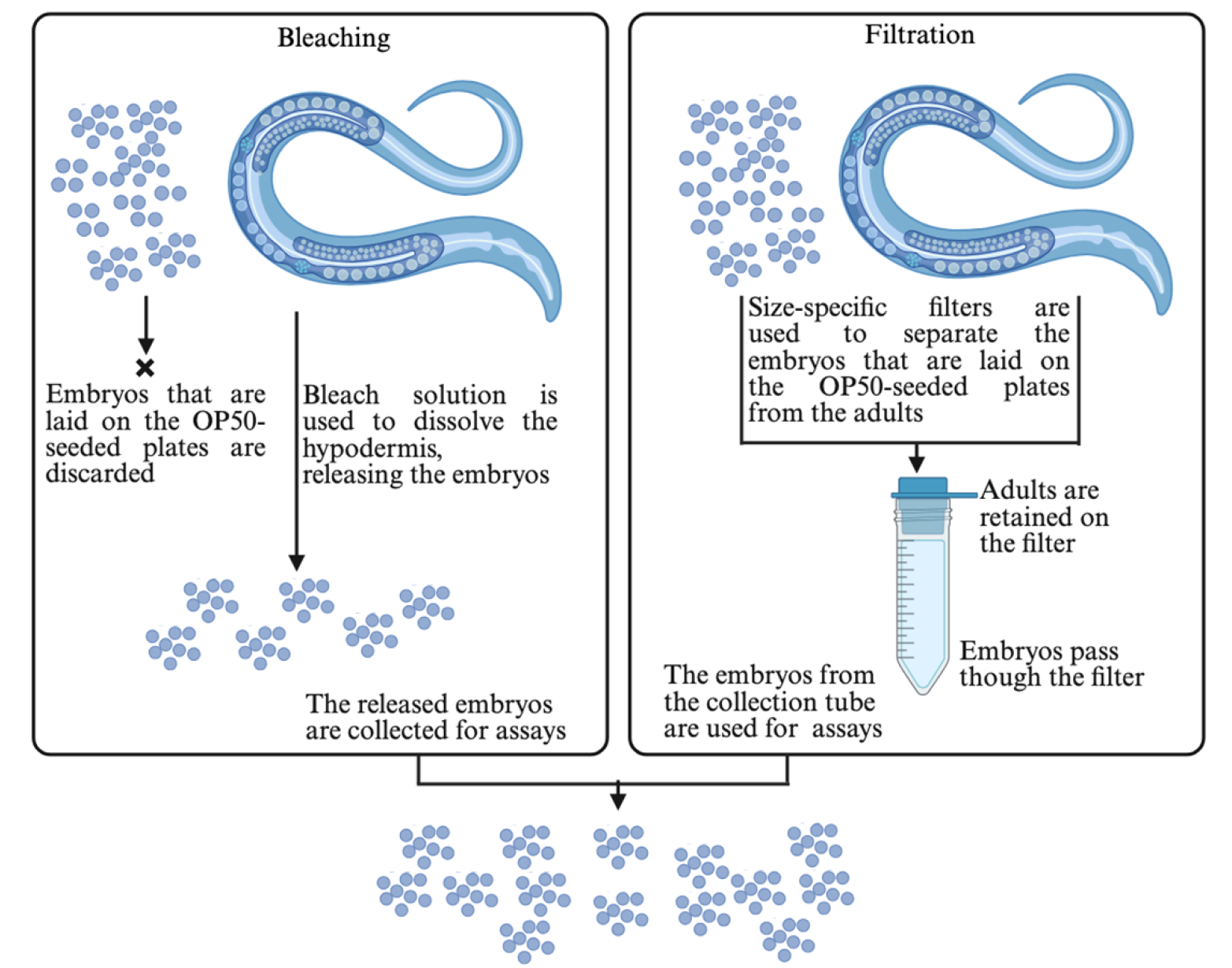
Comparison of bleach and filtration techniques to collect embryos. Bleach synchronization involves dissolving the hypodermis of gravid adults and releasing the embryos. Filtration involves using size-specific filters to separate the embryos from the adults. The figure was created using BioRender.

To overcome the limitations of bleach synchronization, microfluidic devices have been developed that sort animals based on the size differences of each developmental stage (6,7). Such devices enable precise control of parameters such as liquid flow, pressure, and temperature enabling effective sorting (8). However, the major drawback of such devices is their high cost and the requirement to build them, making them inaccessible to many groups.

Mesh filters of different sizes are a cheaper alternative to microfluidic devices. Separation of *C. elegans* stages by size has been done using filters of different mesh sizes that allow specific stages to pass through and some stages to be retained on top of the mesh (Figure 1) (9). Filters with different pore sizes are commercially available. For example, Wilson sieves of 5 µm mesh size were used to separate L1 or L2 populations (10), plate-based nylon mesh filters of 11 µm from Millipore MultiScreen were used to purify the L1 stage (11), pluriSelect filters were used to isolate L4 and young adult stages of *C. elegans* (9), mesh filters have also been used to separate males from hermaphrodites (12), and in-house filters were also constructed with 50 mL conical tubes and monofilament mesh of the required pore size to recover the L4 stage (on 20 µm filters) and day-one adults (on 50 µm filters) (13).

The examples highlight using mesh filters to isolate specific developmental stages of *C. elegans*. However, the small differences in length and width between consecutive larval stages make it difficult to get a 100% pure culture of a specific stage. For example, the difference in width between the L1 and L2 stages, and L2 and L3 stages is approximately 5.3 µm and 5.1 µm respectively (approximate widths of L1, L2, L3, and L4 larvae are 11.7 µm, 17 µm, 22.1 µm, and 29.5 µm, respectively) (6). In contrast, the dimensions of the embryos (50 µm in length and 30 µm in diameter) make it easier to separate from other developmental stages (14). Once isolated, the embryos can be cultured to obtain a synchronous population of L1 larvae that can be grown to a synchronized stage-specific population.

We used filters from pluriSelect to specifically isolate embryos that were laid on an OP50-seeded plate. Based on the widths of the different developmental stages, we conducted sequential filtration using filters of two mesh sizes, 40 µm and 20 µm. The 40 µm filter retained adults and debris. The embryos and larval stages, which are smaller in width, passed through the 40 µm mesh and were collected on the 20 µm mesh. The 20 µm mesh further allowed the smaller larval stages to pass through and the embryos to be retained. Using our technique, we recovered approximately 9,500 embryos from one 6 cm OP50-seeded plate, compared to approximately 5,000 embryos by bleach synchronization, using the *C. elegans* N2 laboratory-adapted strain. To further increase the embryo yield per plate, we incorporated polyethylene glycol (PEG) in the M9 buffer (3% w/v) changing the osmolarity and improving embryo pelleting by three-fold. PEG has been used previously in M9 buffers without adverse effects on *C. elegans* (15). Additionally, PEG has been employed for the encapsulation of various organisms, including *C. elegans*, with no harmful side effects (16). We further showed that the embryos obtained by filtration developed into larvae at a comparable rate to the larvae obtained by bleach synchronization. Furthermore, we found no difference in animal lengths at the L4 stage when embryos were collected using bleach or filtration procedures and exposed to albendazole, an anthelmintic compound known to inhibit animal development (17). In summary, we have developed a protocol to obtain synchronized embryos that bypass the traditional method of bleach synchronization.

## Materials and methods

### Nematode culture maintenance

The strains used for the assays were N2, CB4856, CX11271, ECA36, JU775, and MY16. All strains were from the *Caenorhabditis* Natural Diversity Resource (CaeNDR) (18). Animals were maintained at 20°C for three generations before starting any assay. Modified nematode growth medium (NGMA), which consists of 1% agar and 0.7% agarose was used to grow the strains on 6 cm plates (19). *Escherichia coli* strains OP50, HB101, and X1666 were used as food sources.

### Conical bleach (bleach synchronization) to obtain embryos

Five L4-staged hermaphrodites were transferred to a 6 cm OP50-seeded plate and incubated at 21.5°C for four days. On the fourth day, the animals were washed with M9 buffer (3 g of KH_2_PO_4_, 6 g of Na_2_HPO_4_, and 5 g of NaCl in 1 L Milli-Q water), collected in a 15 mL centrifuge tube, and pelleted by centrifuging at 254 g (Eppendorf 5810R) for one minute. The supernatant was discarded and 6 mL of bleach solution (2 mL of NaOCl and 0.2 g of NaOH in 10 mL Milli-Q water) was added. The tubes were shaken vigorously for four minutes to dissolve the hypodermis. As soon as broken animal bodies were observed under the microscope, the tubes were centrifuged at 254 g (Eppendorf 5810R) for 30 seconds. The supernatant was discarded and approximately 10 mL of M9 buffer was added to the tube and centrifuged at 254 g (Eppendorf 5810R) for one minute. The supernatant was discarded and approximately 10 mL of K medium (51 mM NaCl, 32 mM KCl, 3 mM CaCl_2_, and 3 mM MgSO_4_ in Milli-Q water) was added to the tube and centrifuged at 254 g (Eppendorf 5810R) for one minute. After the final wash, the embryo pellet was resuspended in 3 mL of K medium. The embryo titer was determined by counting the number of embryos in five replicates of 3 µL aliquots.

### Filtration to obtain embryos

The filters and supplementary materials were purchased from pluriSelect, U.S.A. A connector ring (SKU 41-50000) was attached to a 50 mL conical tube. Above the connector ring, a filter stack consisting of a 20 µm filter (green color) (SKU 43-50020) at the bottom and a 40 µm filter (blue color) (SKU 43-50040) at the top was added. A funnel (SKU 42-50000) was attached to the 40 µm filter (Figure 2).

**Figure 2.**
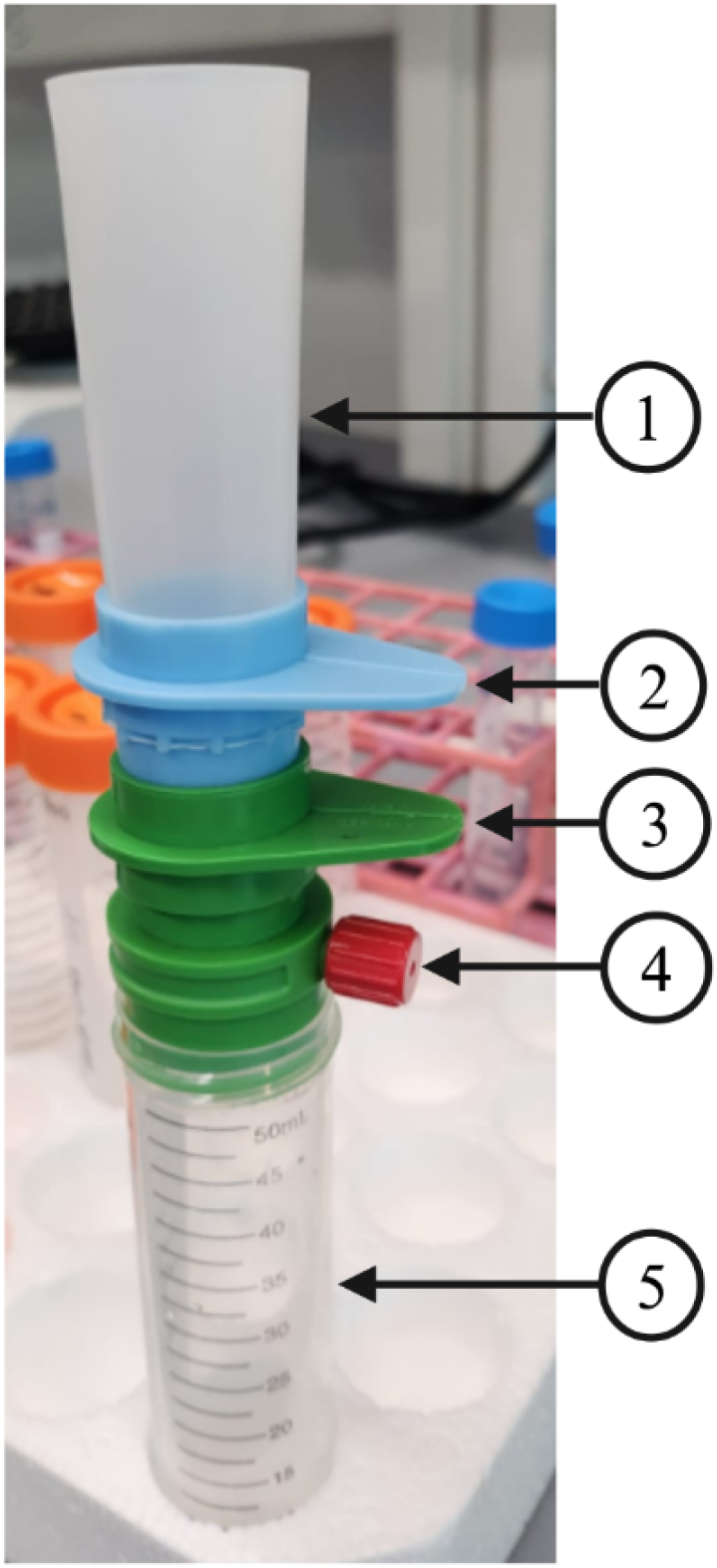
Set up of filtration apparatus. **(1)** A funnel is used to pour the embryo solution. **(2)** 40 µm mesh filter to retain adults. **(3)** 20 µm mesh filter to retain embryos. **(4)** Connector ring to connect to a vacuum to allow the embryo solution to pass through. **(5)** A collection tube is used to collect the flow through, which consists of the majority of larvae.

Embryos were collected using the conical bleach protocol (refer to Conical bleach to obtain embryos in the methods section). After obtaining an average embryo count, the volume of resuspension solution containing 450 embryos was added to 6 cm OP50-seeded plates and incubated at 21.5°C for three days. After three days, the adults were washed from the plates by adding 2 mL M9 buffer, gently swirling the plates, and discarding the solution (Figure 3A). The step was repeated once more. The step ensured that most of the adults and larvae stuck to the OP50 bacteria were washed away (Figure 3B). The embryos retained on the OP50 bacteria lawn were collected by adding 2 mL M9 buffer and gently scraping off the embryos using a rubber policeman (Figure 3C). The M9 buffer containing the embryos was collected in a 15 mL centrifuge tube and passed through the filtration apparatus.

**Figure 3.**
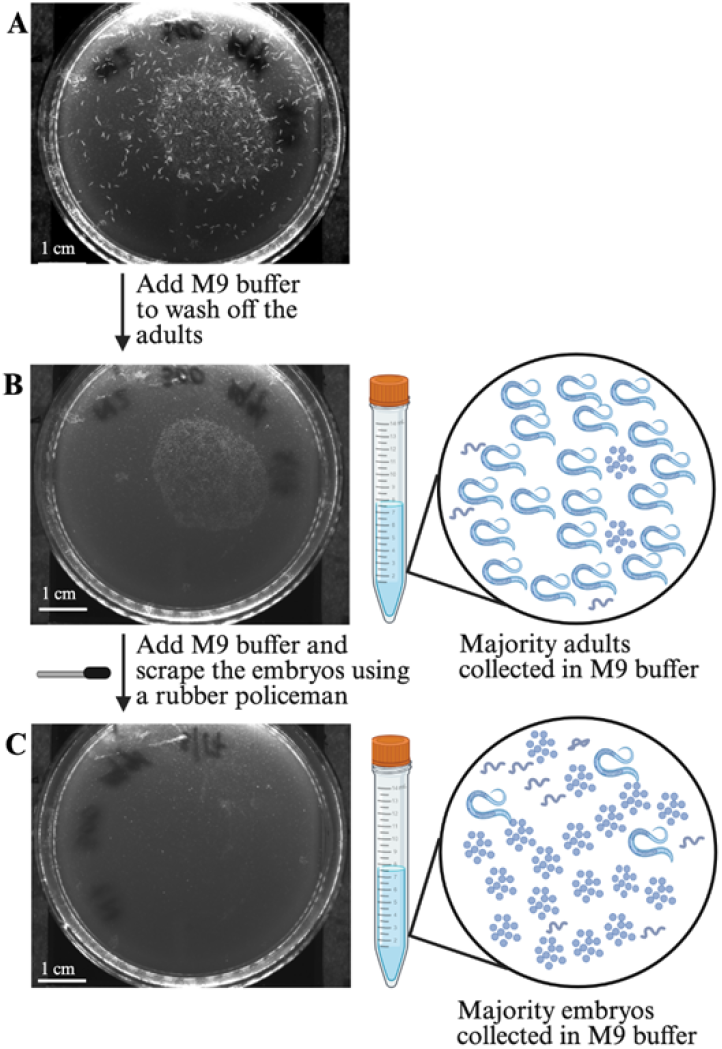
Collection of embryos for filtration. **(A)** A 6 cm OP50-seeded plate with adults and embryos. (**B)** A 6 cm OP50-seeded plate after the adults have been washed away with M9 buffer. **(C)** An OP50-seeded plate after the embryos have been scraped off in M9 buffer with a rubber policeman. The figure was created using BioRender.

The 15 mL tube containing embryos in M9 buffer was inverted 5-6 times to ensure that the embryos were in suspension and added to the funnel of the filtration apparatus (Figures 3C, 4). The connector ring was opened and connected to a vacuum to help the embryo solution pass through the filters. Once all the solution had passed through the filters, the filtration apparatus was disassembled by removing the funnel, connector ring, and 40 µm filter. The 40 µm filter retained the adults and debris, and the embryos and larval stages passed through the filter. The 20 µm filter retained the embryos and a few larvae, and most of the larvae passed through the filter. The 20 µm filter, containing most of the embryos and a few larvae was inverted and added on top of a clean, labeled 50 mL conical tube. The top of the 20 µm filter now faced the inside of the conical tube (Figure 4). 4 mL of 25% bleach solution (1 mL of bleach solution + 3 mL of M9 buffer) was added to the reversed side of the 20 µm filter and incubated for 30 seconds. The addition of 25% bleach killed the few larvae that were retained on the 20 µm filter along with the embryos. After 30 seconds, 6 mL of 3% PEG (PEG3350) (Millipore sigma, Catalog # P4338) in M9 buffer (3 g of PEG in 100 mL of M9 buffer) was added to the filter. After the solution had passed through the filter, the contents of the 50 mL conical tube were transferred to a clean 15 mL centrifuge tube, and centrifuged at 3197 g (Eppendorf 5810R) for one minute. The supernatant was discarded and approximately 10 mL of 3% PEG in M9 buffer was added. The tube was centrifuged at 3197 g (Eppendorf 5810R) for one minute and the supernatant was discarded. The wash step was repeated once more. After the final wash, the embryos were resuspended in 3 mL of K medium. Embryo titer was determined by counting the number of embryos in five 3 µL aliquots.

**Figure 4.**
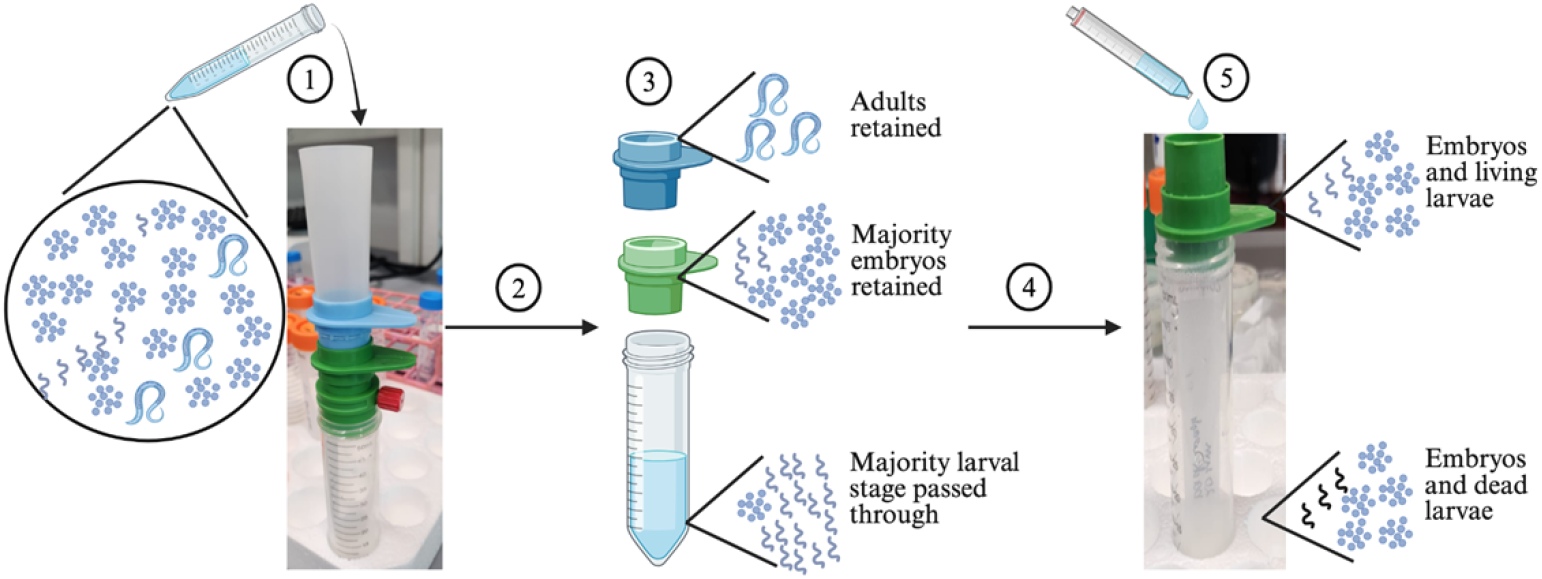
Set up of the filtration apparatus. **(1)** The embryo solution collected in Figure 3C was poured through the filtration apparatus. **(2)** The connector ring was connected to a vacuum to allow the solution to pass. **(3)** After the solution had passed, the filtration apparatus was disassembled. **(4)** The 20 µm filter was inverted and placed on top of a clean 50 mL labeled tube. **(5)** A 25% bleach solution was added to the inverted 20 µm filter for 30 seconds to kill any larvae. The figure was created using BioRender.

### Albendazole stock preparation

Albendazole (Source: Fulka; CAS number: 54965-21-8; Catalog number: A4673-10G) was prepared at a stock concentration of 100 mM in dimethyl sulfoxide (DMSO) and aliquoted in microcentrifuge tubes in 50 µL aliquots that were stored at −20°C. The aliquots were thawed on the day of the assay and discarded after use.

### High-throughput phenotyping assay (HTA)

HTA was carried out as described previously (3). For the developmental analysis, three independent replicates were performed by three experimentalists. For the drug response assay, three independent replicates were performed by one experimentalist. After letting the strains grow for three generations to account for the starvation effects, conical bleach and filtration were carried out to obtain embryos. The embryos were resuspended in K medium to a final concentration of 0.6 embryos per μL, and 50 μL of the embryo solution (containing 30 embryos) was added into each well of a 96-well plate (20). For developmental analysis, 36 wells were prepared for embryos obtained by each protocol (conical bleach and filtration) and by each experimentalist. For the drug assay, 64 wells were prepared for embryos obtained by each protocol (conical bleach and filtration) for all strains. The embryos were incubated at 20°C overnight in K medium with no food and with shaking at 180 rpm to get a synchronized L1 population. The following day, frozen HB101 aliquots at OD_600_ 100 were thawed and diluted to OD_600_ 30 using K medium. HB101 at a concentration of OD_600_ 100 was prepared as previously described (5). 150 μM kanamycin was added to prevent bacterial growth and contamination. For the developmental assay, all 36 wells were treated with control (1% DMSO). For the drug assay, half the wells (32 wells) were treated with 1% DMSO, and the other half were treated with 30 μM albendazole in 1% DMSO. The L1 larvae were allowed to grow for 48 hours at 20°C with shaking at 180 rpm. After 48 hours, the 96-well plates were exposed to 50 mM sodium azide in M9 buffer for 10 minutes to straighten and paralyze the animals. Images of each plate were captured using a 2X objective in a Molecular Devices ImageXpress Nano microscope (Molecular Devices, San Jose, CA) (3). The images were further processed using the CellProfiler, easyXpress, and R studio pipelines (see Data processing and analysis in the methods section) to quantify the development of the animals.

### Data processing and analysis

The raw images were processed using the CellProfiler software (version 4.2.8). The GitHub repository for the pipeline can be found at (https://github.com/AndersenLab/cellprofiler-nf). The processed images generated by CellProfiler were cleaned and further processed using the custom R package easyXpress as previously described (3,21). R studio (version 4.2.1) was used to perform all statistical analysis and figure generation.

## Results and discussion

One of the major challenges of conducting high-throughput *C. elegans* assays is collecting a large number of embryos. Traditionally, bleach synchronization has been used to harvest embryos. However, the process is time-consuming and introduces high variability (3–5). To overcome the challenges of bleach synchronization, we demonstrated the use of commercially available mesh filters to collect a large number of embryos with high purity.

### Optimization of pre-filtration conditions

Our aim was to harvest laid embryos from a 6 cm bacteria-seeded plate. We began this process by plating a fixed number of embryos (P_0_) to ensure that they would develop and lay embryos (G_1_), which could be collected for filtration without the next generation exhausting the bacterial food (strain OP50) on the plate. Different numbers of P_0_ embryos (250, 300, 400, 450, and 500) were added to 6 cm OP50-seeded plates. After incubating the plates at 21.5°C for three days, we observed that adding 450 P_0_ embryos yielded the highest number of G_1_ embryos without food exhaustion. We used 450 P_0_ embryos for all subsequent assays. Next, we sought to determine which *E. coli* bacterial strain enabled most of the G_1_ embryos to remain stuck after P_0_ adults were washed away using M9 buffer. We tested three different *E. coli* bacterial strains -OP50, HB101, and X1666 by plating 450 P_0_ embryos to 6 cm plates seeded with one of the three bacteria strains grown at 21.5°C for three days, after which M9 buffer was added to the plates to wash away the adults. Most G_1_ embryos were washed away along with the P_0_ adults from HB101-seeded plates (Supplementary Figures 1A, B) and X1667-seeded plates (data not shown). In contrast, most of the G_1_ embryos were retained on the plate seeded with OP50 (Supplementary Figures 1C, D). Based on the results, OP50-seeded plates were used for all subsequent assays. In summary, we optimized the pre-filtration conditions by using OP50 as the bacterial food and plating 450 P_0_ embryos for maximum G_1_ embryo yield from one 6 cm OP50-seeded plate. This procedure can be repeated using different numbers of P_0_ embryos if smaller or larger NGMA plates are needed.

### Optimization of filtration conditions

The G_1_ offspring recovered from the 20 µM mesh filter were 94 to 100% (data collected from up to 50 filtrations) embryos with rare L1 larvae contaminants. To get 100% G_1_ embryo cultures, we sought to kill the L1 larvae by exposing the recovered mixed-stage population to a bleach solution. We optimized the bleach concentration and exposure time so that it was sufficient to kill the larvae without having any damaging effects on the G_1_ embryos. We tested three different concentrations of bleach solution (10%, 20%, and 25% in M9 buffer) with varying exposure times (30, 60, 120, and 150 seconds) to identify the optimal condition that killed the larvae (identified by lack of movement) with minimal impact on embryos. Treating the recovered population with 25% bleach solution for 30 seconds killed the larvae with minimal impact on embryonic lethality as measured after 24 hours of recovery (Supplementary Table 1).

During our initial experiments, we used M9 buffer as the wash solution for all subsequent washes. However, embryos might be lost during each M9 buffer wash. To reduce the potential for loss, we altered the osmolarity of the M9 buffer using PEG. We used PEG at four different concentrations (0.5%, 1%, 2%, and 3% w/v) in the M9 buffer and obtained the highest yield with 3% PEG. Consequently, we used 3% PEG in the M9 buffer for all subsequent washes. We recovered two times more embryos from a single 6 cm OP50-seeded plate when 3% PEG was incorporated with M9 buffer in the wash solution (average embryo yield was 12,539 ± 5,727 SD with 3% PEG in M9 buffer compared to an average embryonic yield of 6,821 ± 5,364 SD with M9 buffer) (Supplementary Figure 2A). To ensure that PEG did not affect the hatching of embryos, we assessed the embryonic lethality after 24 hours. The average embryonic lethality of embryos that were washed with 3% PEG was similar to the embryonic lethality of embryos that were washed with M9 buffer (4.5% for 3% PEG in M9 buffer and 3.8% for M9), indicating that the use of PEG does not damage the embryos (Supplementary Figure 2B).

### Filtration yields a higher number of embryos compared to bleach synchronization

To determine the efficacy of embryo filtration, we collected embryos using bleach synchronization and filtration techniques in parallel. We used the same number of 6 cm OP50-seeded plates that had an equivalent number of P_0_ adults. We recovered an average of 4,946 (SD ± 3360) G_1_ embryos by bleach synchronization, compared to an average of 9,474 (SD ± 6764) G_1_ embryos by filtration (Figure 5). To account for variability that might be introduced by different experimentalists performing the protocols, we employed three experimentalists to conduct the filtration and bleach protocols to collect the G_1_ embryos. We did not find a significant difference between any two experimentalists for either of the two protocols (*p* = 0.217 for the bleach synchronization protocol, and *p* = 0.706 for the filtration protocol, Kruskall-Wallis rank-sum test).

**Figure 5.**
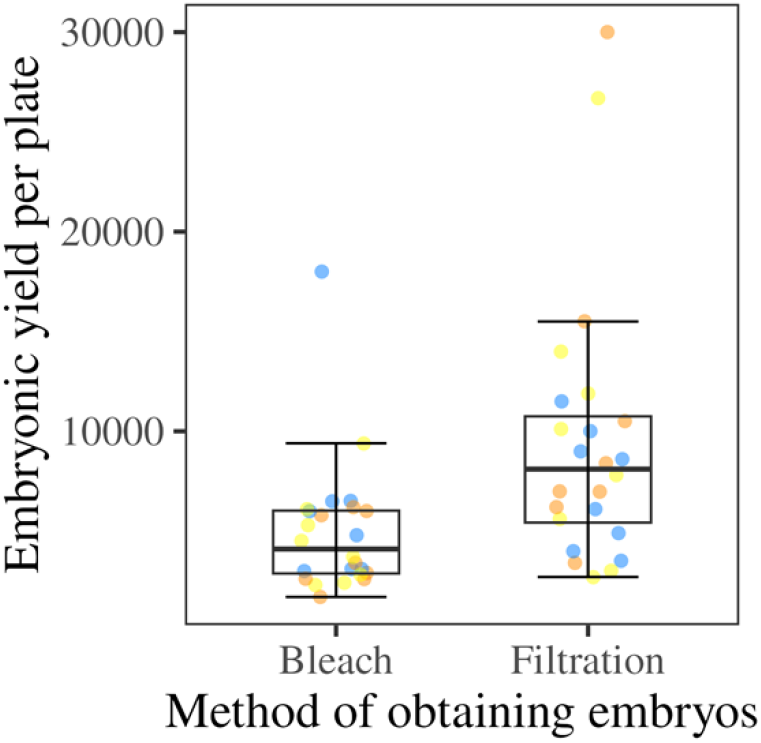
A higher number of G_1_ embryos were recovered from filtration than by bleach synchronization. Box plot showing G_1_ embryonic yield per plate obtained by bleach synchronization or filtration protocols. Median is represented by the solid horizontal line with the 75^th^ and 25^th^ quartiles on the top and bottom of the box, respectively. Whiskers extend from the box to the maximum and minimum values that fall within 1.5 times the interquartile range above and below the 75^th^ and 25^th^ percentiles, respectively. The three different colors of the points indicate three experimentalists. A significantly higher number of G_1_ embryos were collected by filtration (*p* = 0.00064, Wilcoxon rank-sum test). Eight replicates were tested by each experimentalist for each method.

### Embryos obtained by filtration develop comparably to the embryos obtained by bleach

We aimed to understand differences in the development of G_1_ embryos collected by bleach and filtration protocols, so we harvested embryos using bleach synchronization and filtration protocols in parallel and measured the lengths of the animals when they developed to the L4 stage. We used length as an indicator of development because nematodes get longer as they proceed through development. The G_1_ embryos obtained by filtration grew to L4-staged animals having longer or comparable lengths to the lengths of L4-staged animals that developed from G_1_ embryos collected by bleaching, indicating that filtration does not affect larval development (Figure 6).

**Figure 6.**
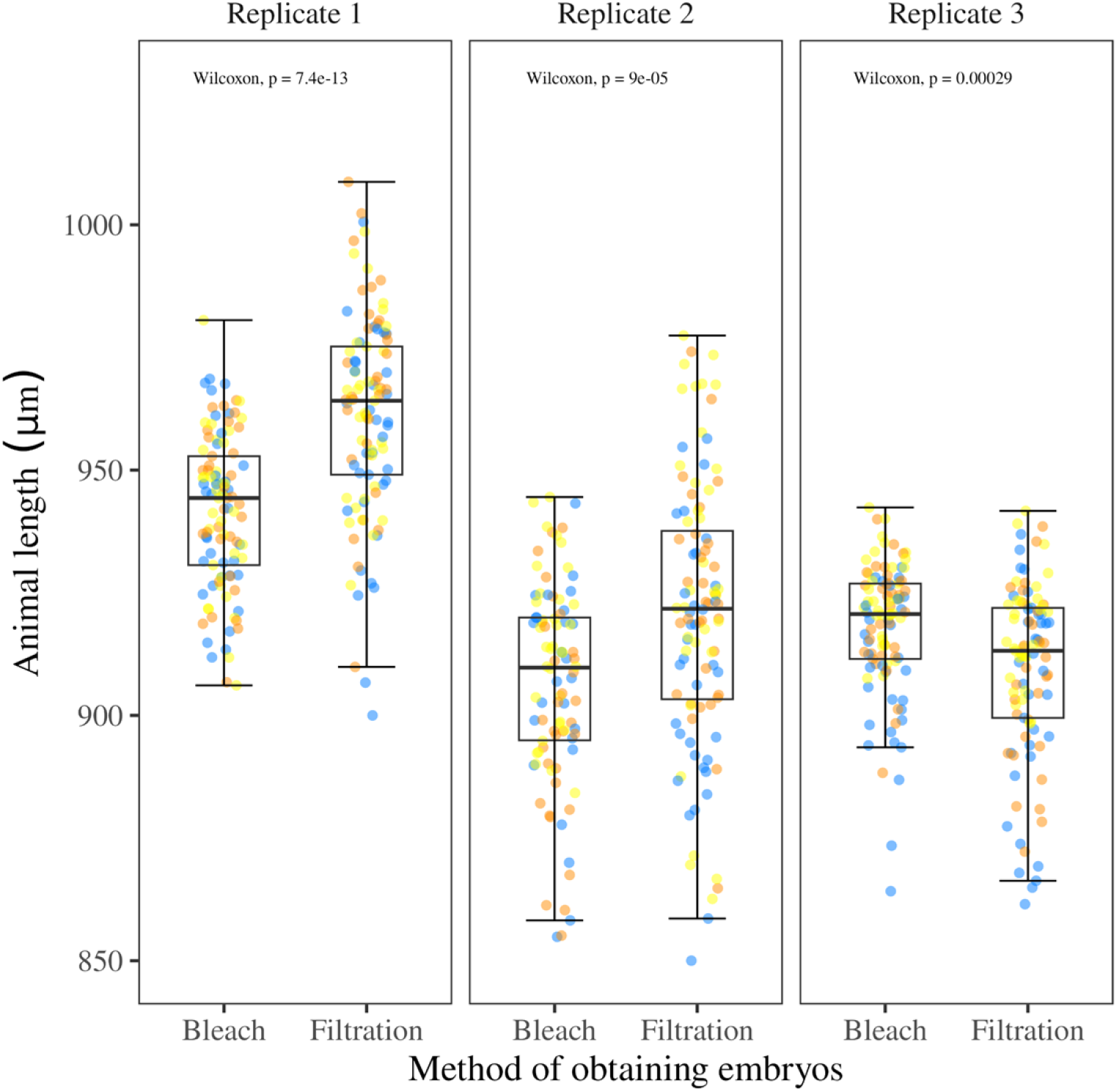
Embryos obtained by filtration developed comparably to the embryos obtained by bleach. The box plot shows the length of L4 animals that developed from G_1_ embryos collected by bleach or filtration protocols. Median is represented by the solid horizontal line with the 75^th^ and 25^th^ quartiles on the top and bottom of the box, respectively. Whiskers extend from the box to the maximum and minimum values that fall within 1.5 times the interquartile range above and below the 75^th^ and 25^th^ percentiles, respectively. Each point is the summarised measurement of an individual well containing between five and 30 animals. Each facet represents a replicate. G_1_ embryos collected by filtration grew into L4 animals that had longer or comparable lengths compared to lengths of L4 animals that developed from G_1_ embryos collected using bleach synchronization, as indicated by *p* values in each facet (Wilcoxon rank-sum test). The assay was carried out independently by three experimentalists (represented by the three different colored points).

### G_1_ embryos collected by filtration develop similarly to the G_1_ embryos obtained by bleach synchronization when exposed to an anthelmintic compound

We aimed to assess whether embryos collected by filtration exhibit a similar response to an anthelmintic drug as the embryos collected by bleach synchronization. We exposed six *C. elegans* strains to albendazole, a benzimidazole drug used to treat parasitic nematode infections in animals and humans (22–24). Benzimidazole drugs act by inhibiting the formation of microtubules, which slows the development of *C. elegans* and can be measured by differences in body length (3,5,25,26). *C. elegans* strains sensitive to albendazole display a shorter body length compared to the resistant strains when exposed to the drug during development (3,17). Natural variation in *C. elegans* response to albendazole has been observed (17). Of the six strains that we selected, four were more sensitive and two were more resistant to albendazole (17). We determined if the results could be replicated using embryos prepared using filtration. For all six strains, the normalized animal lengths for larvae that developed from G_1_ embryos collected by filtration were similar to the normalized animal lengths for larvae that developed from G_1_ embryos recovered by bleach synchronization for both the control (DMSO exposed) and albendazole-exposed conditions (Figure 7).

**Figure 7.**
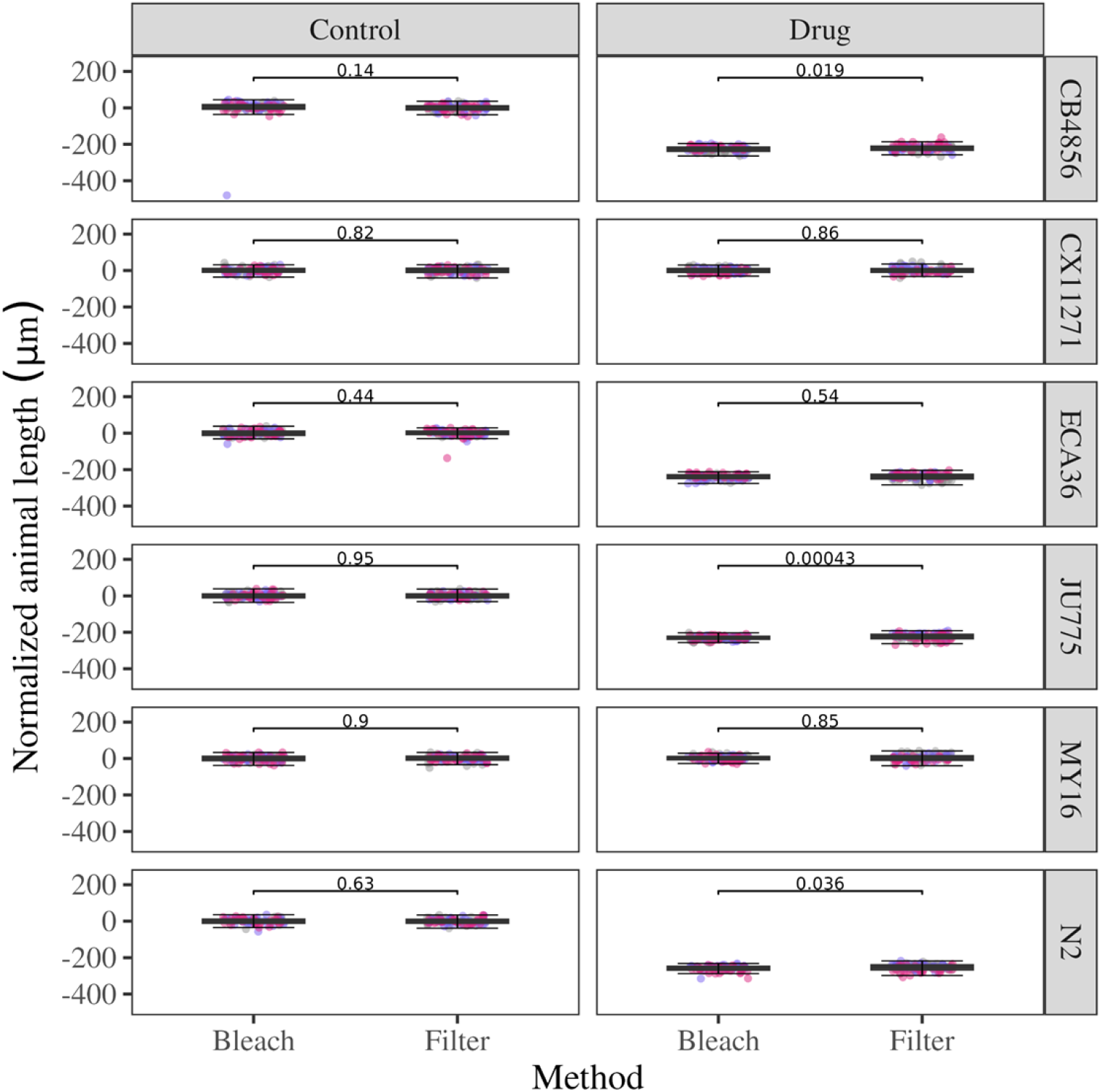
G_1_ embryos collected by filtration developed similarly to the G_1_ embryos obtained by bleach synchronization when exposed to an anthelmintic compound. Box plots showing normalized animal length for larvae developed from G_1_ embryos obtained by bleach synchronization or filtration protocols for six *C. elegans* strains. Median animal lengths of strains exposed to 30 μM albendazole were normalized to the mean of all median animals lengths for the control condition. The median is represented by the solid horizontal line with the 75^th^ and 25^th^ quartiles on the top and bottom of each box, respectively. Whiskers extend from each box to the maximum and minimum values that fall within 1.5 times the interquartile range above and below the 75^th^ and 25^th^ percentiles, respectively. Each point is the summarized measurement of an individual well containing between five and 30 animals. The box plot is faceted by condition (control and drug). No significant difference in the normalized animal length between larvae that developed from G_1_ embryos collected by bleach and filtration for both control and drug conditions was observed, as indicated by the *p*-values (Wilcoxon rank-sum test) in each facet. The assay was replicated three times, as indicated by the colors of the points.

## Conclusion

Bleach synchronization has been routinely used to obtain a synchronized population of embryos. However, because of the harsh chemical nature of bleach, many parameters need to be carefully controlled to ensure the maximum survival of embryos (2). By contrast, filtration takes advantage of the size differences among developmental stages, allowing the separation of embryos without exposing them to chemical stress. We have demonstrated that filtration is a better alternative to bleaching to scale up preparations of staged *C. elegans* embryos in terms of embryonic yield per plate and with no effects on larval development. We optimized our pre-filtration conditions based on the *C. elegans* reference strain N2. Based on the egg-laying rate of N2, we obtained the highest yield of G_1_ embryos by incubating the 450 P_0_ embryos at 21.5°C for three days (27). However, wild strains and mutants have different egg-laying rates and brood sizes, so the number of P_0_ embryos to start the assay might need to be adjusted for other strains. We tested three bacterial strains and found OP50 to be the stickiest to help retain the embryos. Other bacteria might be stickier and should be tested in the future. All our assays were done on 6 cm plates. Higher embryo yield can be obtained by using 10 cm or larger plates. We have applied our technique to harvest embryos from *C. elegans*. However, embryos from other *Caenorhabditis* species whose embryos have similar dimensions to *C. elegans* (e.g., *C. briggsae*) can also be isolated. In summary, our technique provides a robust way of collecting large numbers of *C. elegans* embryos with high purity without the need of subjecting the animals to high concentrations of bleach.

## ACKNOWLEDGEMENTS

We would like to thank members of the Andersen lab for providing their feedback on the manuscript. We also thank *Caenorhabditis* Natural Diversity Resource and *Caenorhabditis* Genetics Center for providing the strains used in the study.

## AUTHOR CONTRIBUTIONS

Nikita S. Jhaveri: Conceptualization, Data curation, Formal analysis, Investigation, Methodology, Validation, Writing - original draft preparation, Writing - review and editing.

Maya K. Mastronardo: Formal analysis, Validation, Writing - review and editing.

J.B. Collins: Formal analysis, Validation, Writing - review and editing.

Erik C. Andersen: Conceptualization, Funding acquisition, Investigation, Methodology, Supervision, Writing - review and editing.

## SUPPORTING INFORMATION

**Supplementary Figure 1**. OP50-seeded plates retained most of the G_1_ embryos after a wash with M9 buffer to remove P_0_ adults.

**Supplementary Figure 2**. The incorporation of 3% PEG in the wash solution improved G_1_ embryonic yield per plate with no effect on embryonic lethality

**Supplementary Table 1**. Optimization of filtration conditions

## DATA AVAILABILITY

Raw data and code are available at the following GitHub repository:

**https://github.com/njhaver4/2025_Filtrationmanuscript**

